# The Rare Plasmid Biosphere: A Hidden Reservoir of Genetic Diversity

**DOI:** 10.1101/2025.07.24.666492

**Authors:** João S. Rebelo, Célia P.F. Domingues, Luís Borda-de-Água, Teresa Nogueira, Francisco Dionisio

## Abstract

Bacterial communities typically display highly uneven abundance patterns, with a few dominant taxa and many low-abundance ones contributing to extensive genetic diversity ^1–4^. Notably, this ‘rare biosphere’ ^5^ includes species performing critical ecological functions, such as biogeochemical cycling and resisting invasions ^6–9^. While bacterial abundance patterns have been extensively studied, the distribution of plasmids-extrachromosomal, self-replicating genetic elements ubiquitous in prokaryotes— remains poorly understood. Using a dataset of 52,909 plasmids from isolates of bacteria and archaea ^10^, we found that their 16,547 Plasmid Taxonomic Units (PTUs) ^11,12^ exhibit a distribution with a fat tail, whether in rank abundance or relative abundance distributions: a few highly prevalent PTUs and many rare. The relative abundance distributions are well described by a Poisson log-normal distribution, consistent with recent findings for species abundance distributions across several taxonomic groups ^13^. The host distribution also presents a heavy tail; however, we show that rare PTUs are not necessarily associated with rare bacterial species, nor are common PTUs exclusively found in common hosts. This indicates that PTUs’ distribution is not a direct consequence of hosts’ distribution. Per plasmid, the host range of rare PTUs is higher than that of common PTUs, at all taxa levels, from species to phyla. Yet, plasmids from common PTUs are more mobile, likely explaining their success. The large group of rare PTUs constitutes a much more diverse reservoir of genetic material than the group of common PTUs. Under appropriate selective pressures, some of these rare plasmids could spread not only by hitchhiking with their hosts but also through horizontal transfer. Therefore, this work opens new paths into plasmid research.

---

Plasmids are pervasive in several bacterial species. For example, *Staphylococcus aureus* strains carry an average of 0.8 distinct plasmids, *Salmonella* 1.7, *Escherichia coli* 2.4, and *Klebsiella* 3.5 ^14,15^. Most plasmids can transfer between different bacterial species, and because they contribute to bacterial virulence, drug resistance, heavy metals resistance, and biofilm formation, they have been extensively studied ^16–18^. In addition, plasmids play a crucial role in bioremediation by carrying genes that encode enzymes and metabolic pathways capable of degrading environmental pollutants and facilitate the metabolism of xenobiotics—synthetic compounds not naturally present in the biosphere. Through horizontal transfer, plasmids can disseminate these traits across bacterial communities, thereby enhancing the bioremediation potential of microbial populations ^19^.

Here, we ask how plasmid diversity is distributed according to the abundance of their taxonomic units. Understanding the distribution of plasmids may be relevant, particularly considering the peculiar distribution of their hosts (mostly bacteria or archaea) within communities. In most ecosystems, bacterial communities are characterized by a few dominant taxa and many low-abundance taxa – the ‘rare biosphere’, representing most species. An interesting characteristic of these rare biospheres ^5^ is their potential for dynamic adaptation, where a vast repertoire of individual cells from low-abundance bacterial taxa can proliferate rapidly through clonal replication when environmental conditions shift to favour their growth ^4,5^.

Answering our question requires a robust plasmid classification, which has been challenging due to the dynamic nature of plasmids, marked by extensive shuffling and recombination ^11^. However, like bacterial chromosomes, plasmids can be clustered, enabling their classification into discrete groups known as Plasmid Taxonomic Units (PTUs) ^11,12^. These clusters meet the criteria of genetic similarity, where members are more similar to each other than to those outside the cluster, and phylogenetic coherence^12^. Therefore, these PTUs are coherent genomic clusters ^20^, similar to the concept of operational taxonomic units (OTUs) in metagenomics, or of bacterial species.

Given the close relationship between plasmids and their prokaryotic hosts, should one expect that a few PTUs dominate, while numerous low-abundance taxonomic units account for most taxa, as is generally observed in the case of bacterial communities ^1,4,21^? In the bacterial world, rare biospheres represent a vast source of genomic innovation ^3,5,21,22^. If a rare plasmid biosphere also exists, it is likely a source of bacterial innovation and may contain ‘hidden’ accessory genes, such as those involved in virulence and antibiotic resistance.

To study the distribution of PTUs, we analysed the RefSeq database containing 52,909 complete plasmids from bacterial and archaeal isolates^10^, spanning 16,547 PTUs.

## Rank-abundance and relative abundance distributions of PTUs

The rank-abundance distribution of the 16,547 PTUs is fat-tailed, revealing a pronounced disparity: a few PTUs are represented by many plasmids, and most PTUs (72%) are represented by only a single plasmid (**Fig. 1a**). In a log-log plot, the distribution of plasmid counts per PTU follows a straight line, suggesting a power-law with a slope close to –0.74 (or –0.91 if we exclude PTUs with a single plasmid) (**Fig. 1b**).

**Fig. 1.**
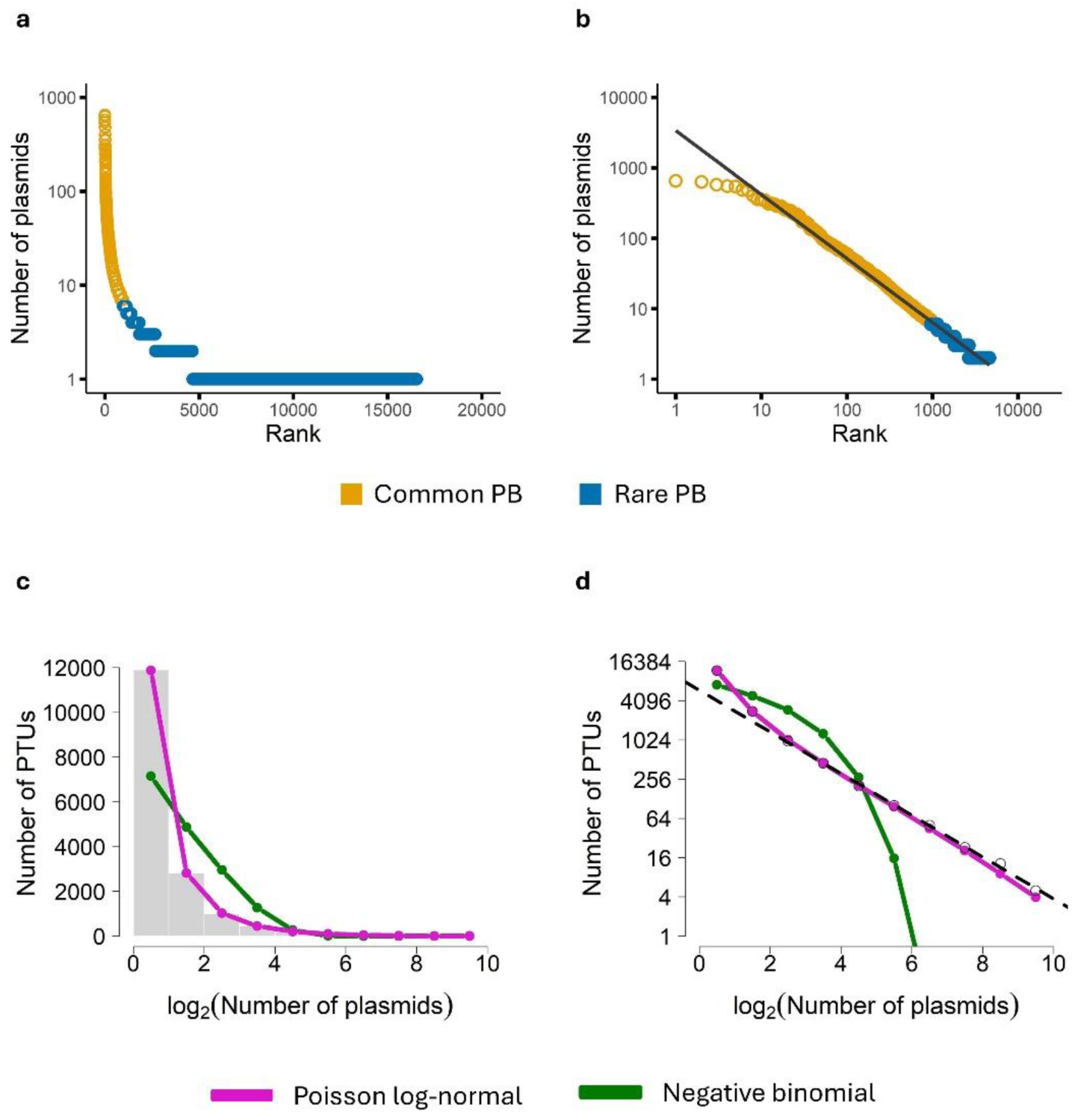
| Rank-abundance and relative abundance distributions of PTUs. **a**, the vertical axis (in log_10_ scale) represents the number of plasmids representing each PTU. The horizontal axis shows the ranking of PTUs, from the most common, with 650 plasmids, to the rarer, with a single plasmid. Yellow: represents the 961 most abundant PTUs, defined as the PTUs with more than 1% of the most abundant PTU (see main text) (Common Plasmid Biosphere); Blue: represents the 15,586 rarer PTUs (Rare Plasmid Biosphere). **b**, the same data as in a, but with horizontal axis in log_10_ scale. The straight line (slope = –0.91 and r^2^ = 0.97) does not consider the plasmids with a single PTU. For this case, the number of plasmids in each rank is given, approximately, by the power-law 3,404.rank^-0.91^. Including all plasmids, slope = –0.74 and r^2^ = 0.89. **c,** shows the PTU abundance distribution (PAD) along with the fits provided by the Poisson log-normal (pink) and the negative binomial (green) distributions. The horizontal axis is in log_2_ scale. **d,** PAD on double logarithmic axes, where we also included a regression line (dashed) fitted to the tail of the distribution, with an estimated slope of approximately –1.063 (excluding the two leftmost points).

Notably, the most abundant PTU contains 650 plasmids, whereas 11,897 PTUs are singletons (i.e., represented by just a single plasmid). How should these numbers be interpreted? If a given PTU has a single representative in the dataset, it means that, among the 52,909 plasmids, no other plasmid in the dataset is similar enough to be included in the same taxonomic unit. We do not know, however, whether this plasmid is very frequent or rare in the habitat (e.g, the gut of an animal) where it was found. We can envisage that this plasmid and the corresponding PTU could be common in a particular habitat, yet no other plasmid of the same taxonomic unit was sampled in the other thousands of habitats^22^.

We also analyse relative abundance distributions, commonly referred to in ecological studies as “species abundance distributions” (SADs), and by analogy, we will refer to them as “PTU abundance distributions” (PADs). These distributions represent, on the x-axis, abundance measured by number of plasmids per PTU, typically shown on a base-2 logarithmic scale (octaves: one plasmid, two or three plasmids, four to seven plasmids, eight to 15 plasmids, *et seq.*), and on the y-axis, the number of PTUs with a given abundance (number of PTUs with one plasmid, with two or three plasmids, and so on). There is an extensive body of ecological literature on fitting SADs ^23,24^. Following recent work ^13^, we fit the PAD using two statistical distributions: the negative binomial (also called Poisson gamma) and the Poisson log-normal distributions, both via maximum likelihood methods ^25^. As can be visually assessed (**Fig. 1c**), the Poisson log-normal distribution provides an excellent fit, in contrast to the negative binomial (see **Supplementary Table S1**, which includes the Akaike information criterion (AIC) values), and **Supplementary Table S2** for the estimated parameters of the distributions). We also plot the PAD in double-logarithmic scales, which further highlights the superior fit of the Poisson log-normal compared to the alternative distribution **(Fig. 1d**).

The fat-tailed distribution implies the existence of many rare PTUs, and the next section will compare these rare PTUs with the common ones. To this aim, we define a PTU as rare if its relative abundance is below a certain threshold. This threshold value implies that the dataset is divided into two ‘plasmid biospheres’ (PBs): a Common PB (the collection of PTUs with many plasmids) and a Rare PB (the collection of PTUs with few plasmids). The concept of rarity has varied thresholds across studies ^3,21^. Because the most abundant PTU has 650 plasmids, we define a PTU as rare if its relative abundance is below 1% – this threshold means that PTUs with fewer than 650×1% = 6.50 plasmids, i.e., with six or fewer plasmids, belong to the Rare PB. Otherwise, they belong to the Common PB. With this threshold, there are 961 PTUs in the Common PB comprising 30,428 plasmids and 15,586 PTUs in the Rare PB comprising 22,481 plasmids (**Figs. 1a and 1b**).

## Are PTUs rare or common because their hosts are also rare or common, respectively?

Plasmids are extra-chromosomal genetic elements, but they can only replicate and remain inside host cells (contrary, for example, to most viruses). Therefore, it would be possible that a rare PTU is rare just because its host species are rare, or that a common PTU is common because its host is common. To test this hypothesis, we looked at the rank-abundance distribution of host species.

Of the 7,715 hosts, 896 have no defined species (they are identified as ‘Genus sp.’). As we intend to perform analyses of the rarity of the host species, and since within a sp. category there may be several species for which we have no information on whether they are common or rare, we have excluded these genomes from the analyses. Therefore, for the host-related analyses, we removed 3,593 plasmids that were identified in hosts with undefined species.

We used two approaches to address the rank-abundance distribution of host species. In the first approach, we consider the number of plasmids sequenced in each of the prokaryotic species. According to this approach, the most frequent host species in the dataset is *E. coli*, harbouring as many as 9,343 plasmids. It is followed by *Klebsiella pneumoniae* (7,977 plasmids), *Salmonella enterica* (2,247 plasmids), *Acinetobacter baumannii* (1,204 plasmids), and 2,359 other prokaryotic species. In the second approach, we count the number of sequenced genomes with plasmids of that species present in the database. According to this approach, the most common species is also *E. coli* (3,068 genomes), followed by *K. pneumoniae* (2,328 genomes), *S. enterica* (1,210 genomes), *Staphylococcus aureus* (807 genomes), and 2,359 other prokaryotic species.

Strikingly, the rank abundance distribution of host species is well fitted by power-laws using both approaches (**Fig. 2a, b**). We then partitioned the hosts’ distribution as we did for plasmids. For both ways of counting hosts, we used a threshold of 1% of the most common host species. Using the first approach to define species frequency (based on the number of plasmids found in a species), the most common host species is *E. coli*, with 9,343 plasmids identified. Therefore, with the threshold of 1%, common hosts species are those with more than 93 plasmids; accordingly, there are 58 common and 2,305 rare host species. For the second approach, a threshold of 1% means that a species is common if the database contains at least 30 sequenced genomes, because the most frequent host species in the database is *E. coli* with 3,068 genomes with plasmids. Accordingly, there are 68 common and 2,295 rare host species.

**Fig. 2.**
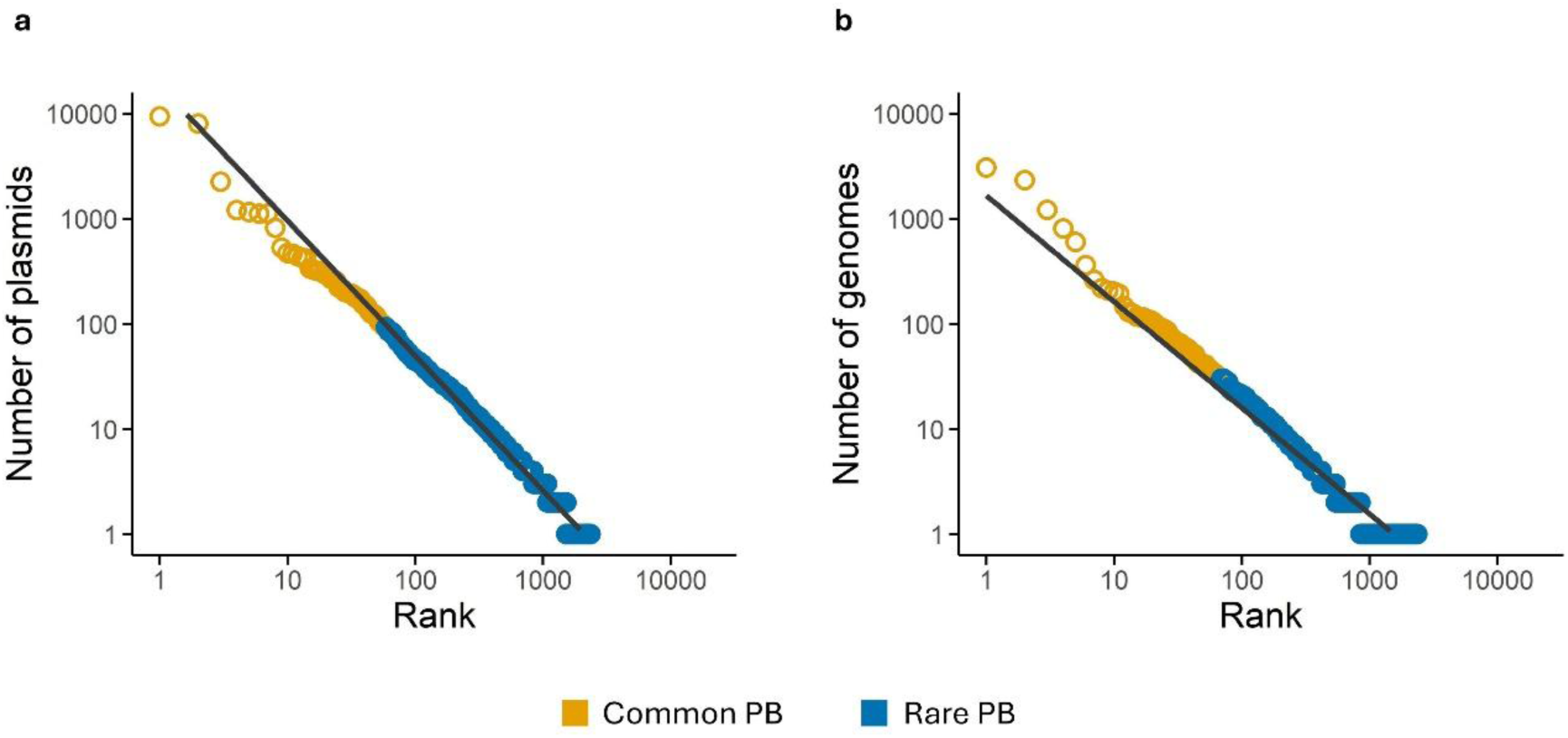
| Rank-abundance of host species. **a**, Rank of host species according to the number of plasmids found in the species (first approach – see main text). The first point on the left (rank = 1) represents *E. coli*, which has 9,343 plasmids. Both scales are in Log_10_. The straight line represents the power-law with slope –1.28, R^2^ = 0.98. **b,** Rank of host species according to the number of genomes with plasmids of each species in the database (second approach – see main text). The first point (rank = 1) on the left represents the 3,068 *E. coli* genomes with plasmids. Both scales are in Log_10_. The straight line represents the power-law with slope –1.01, R^2^ = 0.94.

We next asked whether rare host species preferentially carry plasmids from rare PTUs and common host species carry plasmids from common PTUs. If plasmids were randomly distributed between hosts, 57% of them would inhabit a host whose biosphere category (rare or common) does not match their own category. Using the first classification approach, we identified 13,534 plasmids in rare hosts: 3,035 from the Common PB and 10,499 from the Rare PB. Of the 35,782 plasmids in common hosts, 26,887 belong to the Common PB and 8,895 to the Rare PB. Thus, 3,035 + 8,895 = 11,930 plasmids (24.2% of the total) do not match the biosphere category of their hosts. With the second approach, 13,529 plasmids were associated with rare hosts (3,218 from the Common PB; 10,311 from the Rare PB) and 35,787 with common hosts (26,704 from the Common PB; 9,083 from the Rare PB), yielding 3,218 + 9,083 = 12,301 mismatched plasmids, or 24.9%. Hence, in both approaches, roughly one-quarter of plasmids reside in hosts of the opposite biosphere category, far fewer than the 57% expected under random assignment, but well above a perfect correspondence (0%).

The above assessment overlooks what happens within each species. We examined the association between the total number of plasmids found in a species (first approach) or between the total number of sequenced genomes of a host species (second approach) and the percentage of plasmids of the Rare PB in that species. If plasmids from the Rare PB are associated with rarer species, both correlations should be negative: most plasmids found in rare species would belong to the Rare PB, while the percentage of plasmids from the Rare PB would be low in more common species. The correlation is indeed negative, but the association is moderate only, and the widely dispersed data points suggest that other factors may play a more significant role in determining the relationship between PTU and host species abundances (**Supplementary Fig. 1**).

Both approaches for defining rare and common host species yielded essentially the same pattern – qualitatively identical and quantitatively very similar. Therefore, the following sections present only the results based on the first approach.

Because prokaryote species may host plasmids both from the Common and Rare PBs, we conducted a binomial test for each common species to assess whether the proportion of common plasmids, (number of plasmids from the Common PB) / (total number of plasmids), is significantly different from the proportion of common plasmids in the sample, namely 30428/52909 = 0.58. Among the 58 common species, 18 host more plasmids from the Rare PB than expected (binomial test, *p* < 0.05), and 11 common species do not have more plasmids of one plasmid biosphere than expected (binomial test, *p* > 0.05). The other 29 common species contain more plasmids from the Common PB than expected (binomial test, p < 0.05). Yet, even among these 29 species, there are many plasmids belonging to the Rare PB (**Supplementary Fig. 2**). We then performed a similar analysis with the rare species (hence with 93 plasmids or fewer, following the first approach). There are 2,305 species in these conditions: 21 contain more plasmids of the Common PB than expected (binomial test, *p <* 0.05), and 1,686 do not contain more plasmids of any biosphere than expected (binomial test, *p >* 0.05). In 598 rare species, there are more plasmids from the Rare PB than expected (binomial test, *p <* 0.05), but even in these species, there are plasmids of the Common PB. In the particular case of PTUs with just one plasmid (9,692 singletons), 6,307 (approximately 2/3 of singletons) were found in rare host species and 3,385 (∼ 1/3) in common host species. Notably, every host species classified as common harboured at least three plasmids assigned to the Rare PB. On the other hand, 59 rare host species harbour plasmids only from the Common PB. Overall, these analyses suggest that plasmids of common PTUs are not common just because they were found in common host species, as we found so many plasmids of the Rare PB in the common hosts and plasmids of the Common PB in rare host species. These analyses indicate that plasmids from the Rare PB tend to be associated with rare prokaryotic species, but this trend is only moderate, as so many plasmids of the Rare PB are found in common species (**Supplementary Figs. 1-3**).

## The Rare Plasmid Biosphere is Richer, Has More Distinct Virulence Genes and Fewer Antibiotic Resistance Genes

Because each common PTU contains more plasmids than every rare PTU, PTUs in the Common PB can be viewed as the more “successful” lineages. To test whether this success reflects larger plasmid size, and hence a greater gene repertoire, we compared plasmid lengths (in base pairs) between the Common and Rare PBs. The size of plasmids does not differ between the two groups, indicating that the greater abundance of common PTUs cannot be explained by longer genomes **(Supplementary Fig. 4a)**. Moreover, there are fewer coding DNA sequences (CDS) clusters in the Rare PB than in the Common PB, but the strength of the association is irrelevant, so the statistical significance is driven by a large sample size **(Supplementary Fig. 4b)**. Therefore, data does not support the hypothesis that plasmids from the Common PB are more abundant because they contain more CDS clusters.

The two biospheres together account for 875,346 CDS clusters, but the Common PB has only 76,491 CDS clusters (8.74%) while the Rare PB has 826,373 (94.41%), a significant difference (the combined percentages exceed 100% because some CDS clusters are present in both plasmid biospheres). Another way to see the biological significance of this difference is that, while 48,973 CDS clusters are specific to the Common PB (i.e., absent from the Rare PB), 798,855 CDS clusters are only found in the Rare PB. The difference in the number of CDS clusters between the two plasmid biospheres is significant. Moreover, although there are fewer plasmids in the Rare PB, they have significantly more CDS clusters than the Common PB (**Fig. 3**).

**Fig. 3.**
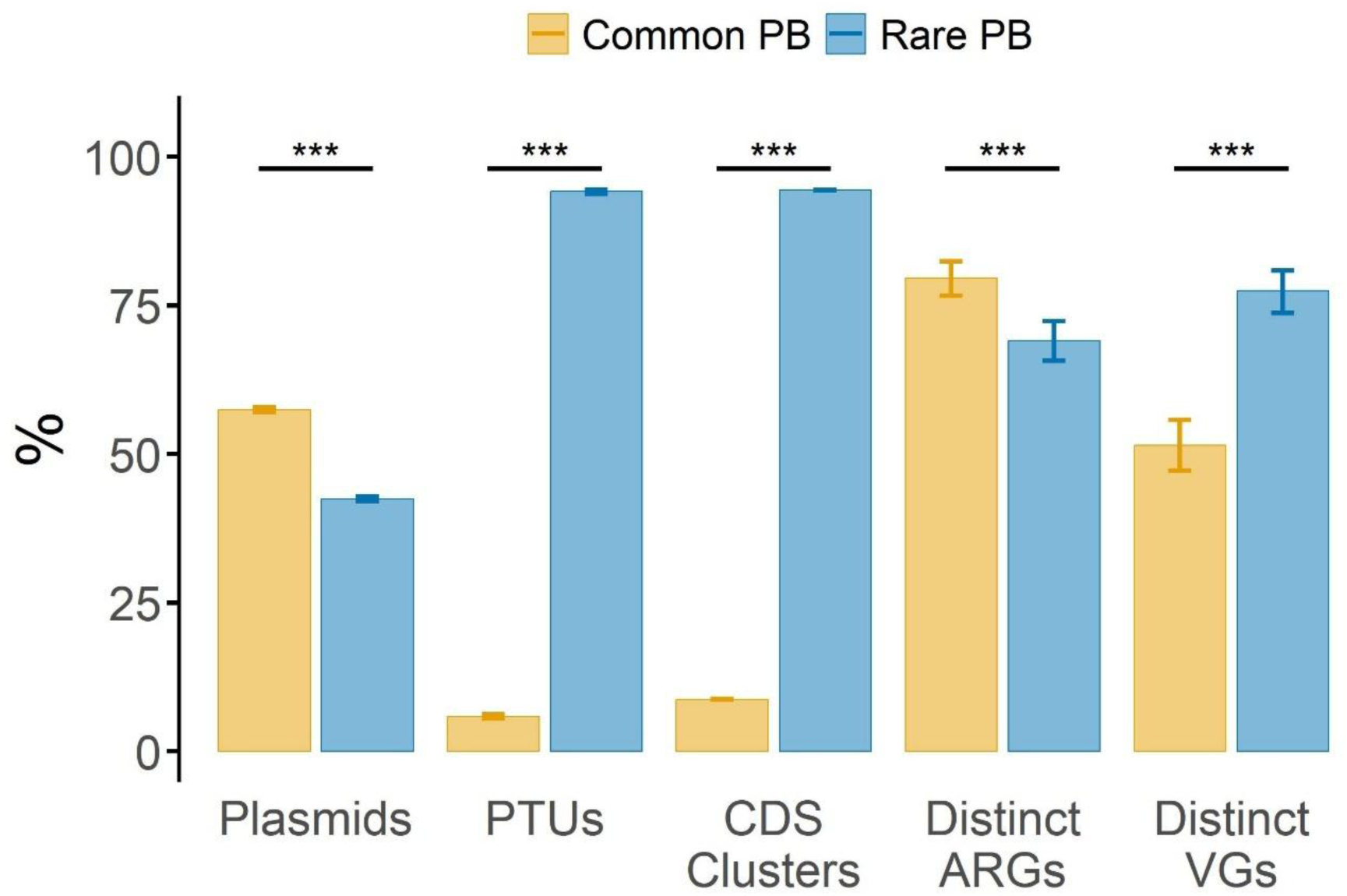
| Characteristics of the Common and Rare Plasmid Biospheres. The Rare PB includes fewer plasmids than the Common PB (*χ*^2^(1) = 2387, *p <* 2.2×10^−16^, Cramér’s V = 0.150, CI_95%_: [0.145, 0.157]). The number of PTUs is higher in the Rare PB than in the Common PB (*χ*^2^(1) = 25849, *p <* 2.2×10^−16^, Cramér’s V = 0.884, CI_95%_: [0.879, 0.889]). The number of CDS clusters is also higher in the Rare PB than in the Common PB (χ^2^ (1) = 1286069, *p <* 2.2×10^−16^; Cramér’s V = 0.857, CI_95%_: [0.856, 0.858]). The number of distinct VGs is higher in the Rare PB than in the Common PB (χ^2^ (1) = 80, *p <* 2.2×10^−16^, Cramér’s *V* = 0.272, CI_95%_: [0.208, 0.332]). The number of distinct ARGs is higher in the Common PB than in the Rare PB *(χ*^2^ (1) = 22, *p* = 3.5×10^−6^, Cramér’s V = 0.120, CI_95%_: [0.069, 0.167]). Bars on columns indicate the CI_95%_.

Among the 4,814 plasmids containing virulence genes (VGs), there are only 546 distinct VGs, so some of them are found more than once across plasmids. The Common PB contains 4,053 plasmids encoding VGs but only 281 distinct VGs. The Rare PB, on the other hand, includes only 761 plasmids with VGs, but 423 distinct VGs. Therefore, only 51.5% (CI_95%_: [47.2%, 55.7%]) of distinct VGs are in the Common PB, whereas 77.5% (CI_95%_: [73.7%, 80.9%]), are in the Rare PB, with an odds ratio of 0.31 (CI_95%_: [0.24, 0.40]), indicating a significantly higher prevalence of these genes in the Rare PB. Hence, the diversity of virulence genes is higher in the Rare PB than in the Common PB (**Fig. 3**). The biological significance of this difference becomes particularly evident when considering the following: of the 546 distinct VGs identified, the Rare PB includes 265 that are absent from the Common PB. As such, failing to detect the Rare PB would result in overlooking 48.5% of the total VGs. In contrast, only 123 VGs (22.5%) found in the Common PB are not present in the Rare PB.

The Virulence Factor Database (VFDB) classifies virulence genes into 14 categories ^26^. One category – post-translational modification – was absent from all plasmids in the dataset. However, 12 categories were identified in both plasmid biospheres (**Supplementary Fig. 5a**). No category was found exclusively in the Common PB, but the stress survival category was found exclusively in the Rare PB, represented by the *sodCI* gene. This gene, typically present in the chromosome of *Salmonella enterica* serovar Typhimurium, was detected in two plasmids. The *sodCI* gene encodes the superoxide dismutase SodCI, which enhances the virulence of these bacteria by protecting them against macrophages^27^.

Concerning antibiotic resistance genes (ARGs), we found 79.6% (CI_95%_: [76.6%, 82.4%]) of the 761 different ARGs in the Common PB, whereas 69.1% (CI_95%_: [65.7%, 72.4%]) of the ARGs were present in the Rare PB, with an odds ratio of 1.75 (CI_95%_: [1.38, 2.21]), indicating that the odds of ARG presence in the Common PB were higher than those in the Rare PB (**Fig. 3**). The dataset contains 14,718 plasmids with 761 distinct ARGs, of which 371 are present in both plasmid biospheres, 155 are only present in the Rare PB, and 235 in the Common PB. Therefore, although the Common PB is richer in ARGs than the Rare PB, overlooking the Rare PB would mean ignoring the presence of the 155 ARGs specific to this plasmid biosphere.

According to AMRFinderPlus, plasmids in the dataset contain genes conferring resistance to 28 distinct antibiotic categories (**Supplementary Fig. 5b**) ^28^. Of these, 27 are found in both plasmid biospheres, while one category—Nitroimidazole—is exclusive to the Rare PB. This category includes the genes *nimA* and *nimD* found in plasmids from *Bacteroides fragilis*, and *nimE* found in plasmids from *Bacteroides fragilis*, *Bacteroides* sp. PHL 2737, and *Bacteroides xylanisolvens*.

## PTUs of the Rare PB Spans Through More Host Species, and, *per* Plasmid, have Broader Host-Range

Plasmids of the Rare PB represent only 42.5% of the 52,909 plasmids analysed in this study. Yet, they span more taxa than the plasmids from the Common PB (**Table 1**). For example, there are 27 hosts’ phyla with plasmids belonging to the Rare PB, but just 11 hosts’ phyla with plasmids of the Common PB. (**Supplementary Fig. 6**). Moreover, there are no hosts’ phyla harboring plasmids only of the Common PB, but there are 16 hosts’ phyla with plasmids only of the Rare PB. This pattern repeats at all taxa levels until the species level (**Table 1**). At this level, there are 2,304 host species with plasmids of the Rare PB, but only 465 with plasmids of the Common PB. Also, 744 host species harbour plasmids only of the Rare PB but only eight harbour plasmids only of the Common PB (**Table 1**; as explained before, we removed from the analysis at the species level the plasmids identified in genomes with unidentified species: 3,593 plasmids, 87 from archaea, and 3,506 from bacteria).

**Table 1:**
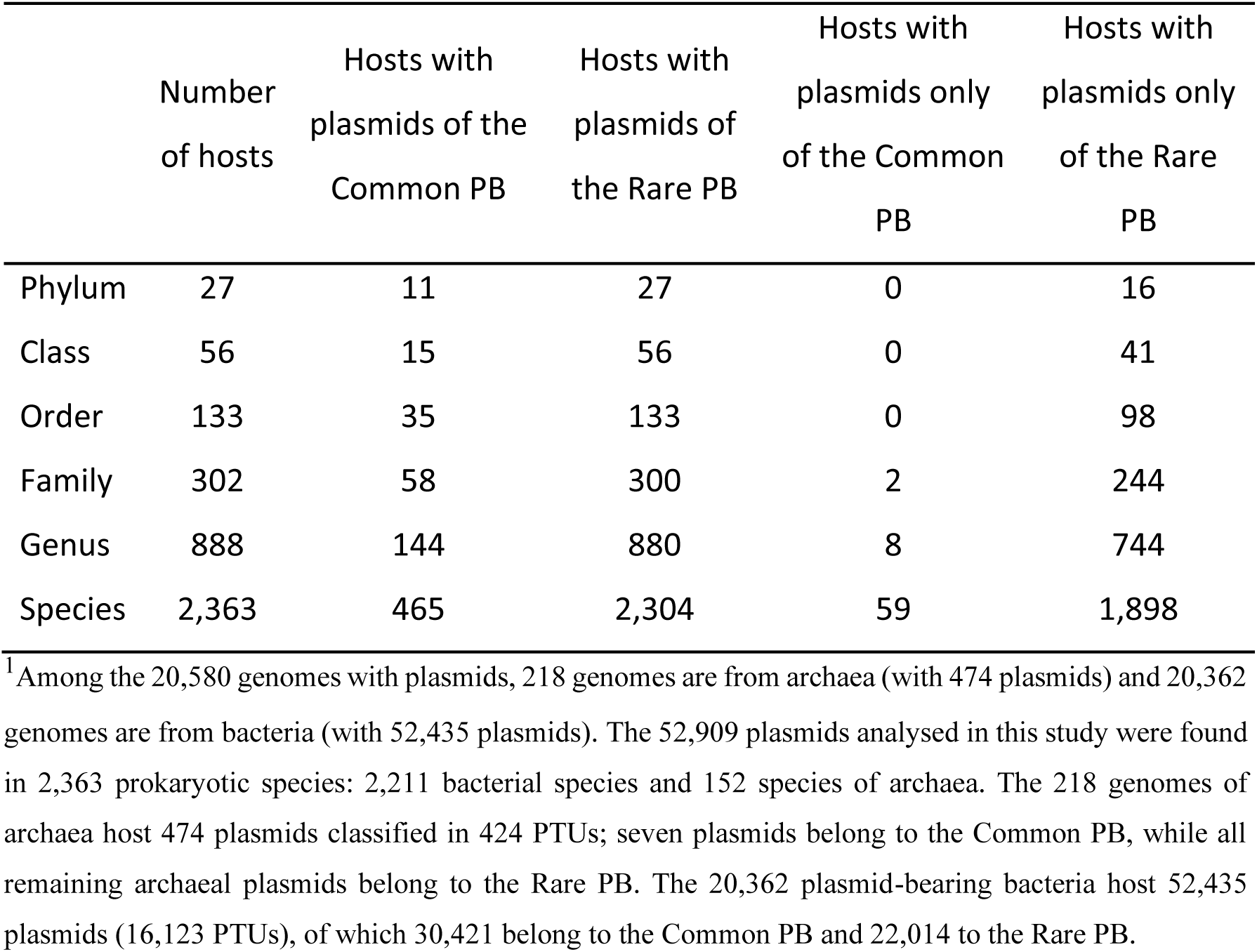
Taxonomy of host species harbouring plasmids.^1^.

|  | Number<br>of hosts | Hosts with<br>plasmids of the<br>Common PB | Hosts with<br>plasmids of<br>the Rare PB | Hosts with<br>plasmids only<br>of the Common<br>PB | Hosts with<br>plasmids only<br>of the Rare<br>PB |
| --- | --- | --- | --- | --- | --- |
| Phylum | 27 | 11 | 27 | 0 | 16 |
| Class | 56 | 15 | 56 | 0 | 41 |
| Order | 133 | 35 | 133 | 0 | 98 |
| Family | 302 | 58 | 300 | 2 | 244 |
| Genus | 888 | 144 | 880 | 8 | 744 |
| Species | 2,363 | 465 | 2,304 | 59 | 1,898 |

<sup>1</sup> Among the 20,580 genomes with plasmids, 218 genomes are from archaea (with 474 plasmids) and 20,362 genomes are from bacteria (with 52,435 plasmids). The 52,909 plasmids analysed in this study were found in 2,363 prokaryotic species: 2,211 bacterial species and 152 species of archaea. The 218 genomes of archaea host 474 plasmids classified in 424 PTUs; seven plasmids belong to the Common PB, while all remaining archaeal plasmids belong to the Rare PB. The 20,362 plasmid-bearing bacteria host 52,435 plasmids (16,123 PTUs), of which 30,421 belong to the Common PB and 22,014 to the Rare PB.

Recently, Redondo-Salvo et al. classified ^12^ host-ranges of PTUs into a scale of six grades, ranging from I (restricted to a single host species) to VI (able to colonize different phyla). For example, the 32 host species harbouring the 637 plasmids (more 13 plasmids from five unidentified species) of the largest PTU belong to a single family (Enterobacteriaceae) and just 12 genera (*Citrobacter*, *Cronobacter*, *Enterobacter*, *Escherichia*, *Klebsiella*, *Kluyvera*, *Leclercia*, *Phytobacter*, *Pseudescherichia*, *Raoultella*, *Salmonella*, *Shigella*). Although this is the largest PTU, it is present in different genera within a single family. Therefore, it is of ‘intermediate’ host-range, i.e., grade III. In the Common PB, a PTU comprising 24 plasmids is found across three phyla: Campylobacterota, Bacillota (Firmicutes), and Actinomycetota. This is a case of a very broad host-range (grade VI). However, the PTU found in more phyla is present in four phyla (Bacillota, Verrucomicrobiota, Actinomycetota and Pseudomonadota) despite belonging to the Rare PB. Therefore, even rare PTUs may be distributed among very distinct taxonomic groups. This was unexpected because, by definition, a PTU belongs to the Rare PB if it has up to six plasmids in the dataset. This specific PTU present in four phyla has only four plasmids. On the other extreme, 325 Common PTUs are in a unique host species, hence very narrow host-range plasmids (grade I) (**Supplementary Fig. 7**).

Next, we analyse and compare the host-range of Common and Rare PTUs in a systematic way. It is likely that common PTUs are present in more taxa (e.g., in more species) than rare PTUs simply because common PTUs are, according to our definition, those having more representative plasmids in the dataset (therefore with more opportunities to be found in more taxa). To rule out this confounding factor concerning host-range, we calculated the number of hosts’ species per number of plasmids of each PTU. This ratio is significantly higher (Wilcoxon signed-rank, p < 0.05) for the PTUs of the Rare PB than of the Common PB **(Fig. 4a).** Since thousands of PTUs contain only one plasmid (singletons), we conducted a similar comparison excluding these PTUs from the analysis. Again, the diversity of hosts’ species per plasmid is higher in the Rare PB than in the Common PB **(Fig. 4b)**. These ratios may be too low in the bigger PTUs. For example, the hosts’ species/plasmid ratio for a PTU with 30 plasmids present in 5 species (ratio = 5/30) is lower than for a PTU with 5 plasmids in 5 species (ratio = 5/5). To avoid these biases, we compared the success of PTUs with seven plasmids (belonging to the Common PB) with those with six plasmids (belonging to the Rare PB). Again, for this case, the PTUs of the Rare PB are present in more host species, on average, than the PTUs of the Common PB **(Fig. 4c).** A similar analysis for genus, family, order, class and phylum has shown that this ratio is consistently higher in the Rare PB than in the Common PB (Wilcoxon signed-rank test *p* < 0.05, large effect sizes for all cases) (**Figs. 4d, 4e, and 4f, and Supplementary Fig. 8** show these analyses for the genus, family, order, and class levels). Therefore, other factors must be responsible for the success of Common PTUs.

**Fig. 4.**
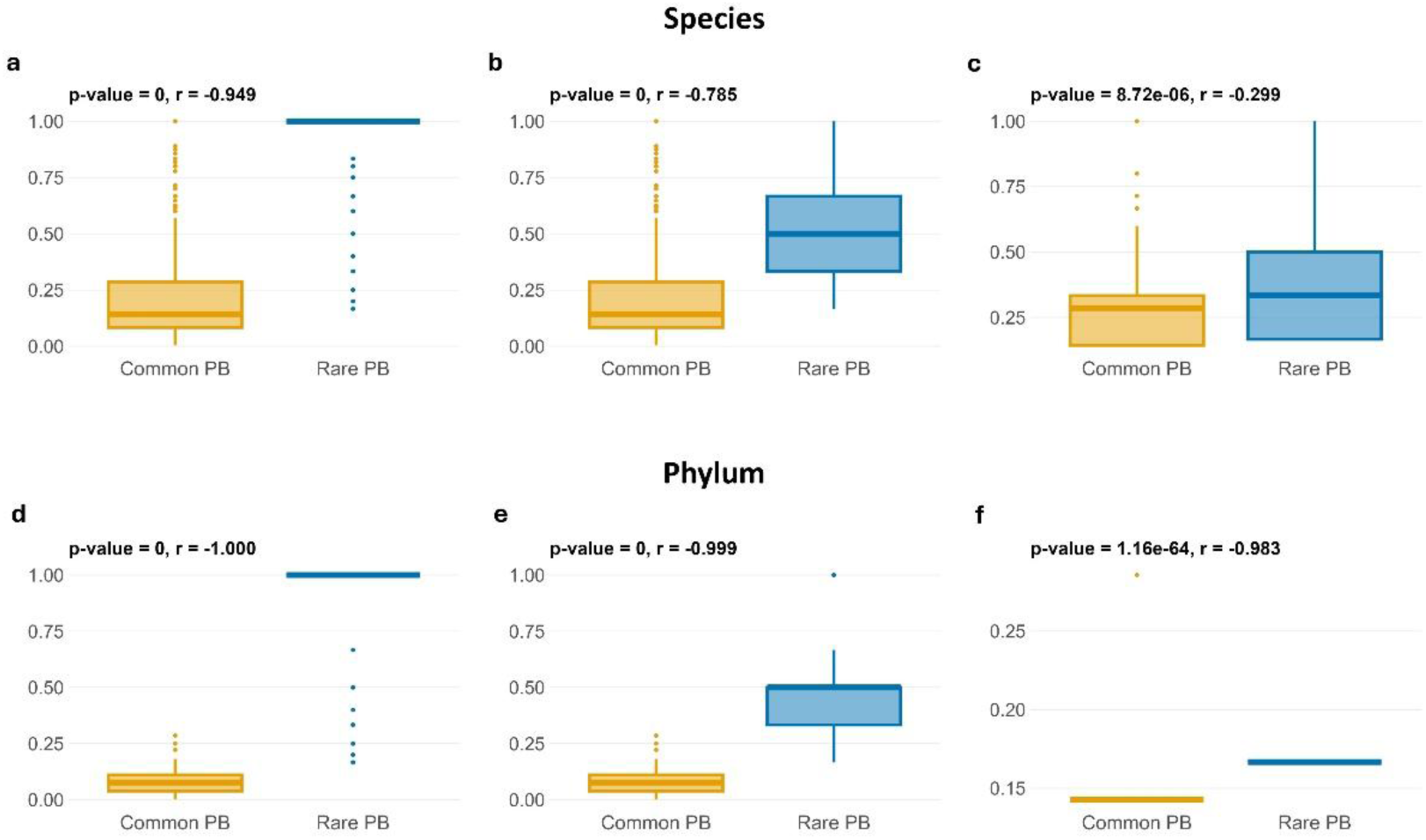
| The relative host-range of PTUs of the Rare and Common PB. The relative success of each PTU, measured as the number of hosts’ taxa per number of plasmids in that PTU. From left to right: all plasmids, not including singletons, including only PTUs with seven (Common) or six (Rare) plasmids, respectively. **a, b, c**, species level. **d, e, f**, phylum level. Host-range is higher for the PTUs of the Rare PB than that of the Common PB for both levels (Wilcoxon signed-rank test *p <* 0.05, large effect size in all cases, except for the species level, where the effect size is moderate, r = – 0.299, when comparing PTUs with six versus seven plasmids).

## Plasmid mobility is higher among plasmids of the Common PB

With the success of the most common PTUs still unexplained, we looked for their transfer ability. Plasmid mobility falls into four main classes. Conjugative plasmids (pCONJ) encode all the necessary genes for self-transfer by conjugation, namely a relaxase, an origin of transfer (*ori*T) and a type IV secretion system. Mobilizable plasmids (pMOB) retain only part of this machinery—typically a relaxase and an origin of transfer, *ori*T—and “hitch-hike” on the conjugation systems of co-resident plasmids. Some plasmids, pOriT, encode just *ori*T; these plasmids need to “hitch-hike” on the conjugation system and the relaxase of other plasmids in the same cell. Finally, some plasmids are non-transmissible (pNT), lacking all transfer genes and being inherited only through cell division or hitchhiking mechanisms such as conduction (co-integration in a transferable plasmid), transduction (with the help of a phage), or transformation ^14,29–33^. The proportions of each class in the Common and Rare PBs tend to be significantly different from the expected values if they were randomly distributed through the PBs (Table 2).

**Table 2.** | Plasmids’ transfer ability in the Common and Rare PBs.

|  |  | N | Obs (%) | CI<sub>95%</sub> | E (%)<sup>1</sup> | CI<sub>95%</sub> |
| --- | --- | --- | --- | --- | --- | --- |
| Common PB | pCONJ | 10928 | 35.9 | [35.4, 36.4] | 28.5 | [28.0, 29.0] |
|  | pMOB | 6438 | 21.2 | [20.7, 21.6] | 22.5 | [22.0, 23.0] |
|  | pOriT | 5871 | 19.3 | [18.8, 19.7] | 16.5 | [16.1, 16.9] |
|  | pNT | 7191 | 23.6 | [23.1, 24.1] | 32.5 | [31.9, 33.0] |
| Rare PB | pCONJ | 4174 | 18.6 | [18.1, 19.1] | 28.5 | [27.9, 29.1] |
|  | pMOB | 5460 | 24.3 | [23.7, 24.8] | 22.5 | [21.9, 23.0] |
|  | pOriT | 2860 | 12.7 | [12.3, 13.2] | 16.5 | [16.0, 17.0] |
|  | pNT | 9987 | 44.4 | [43.8, 45.1] | 32.5 | [31.8, 33.1] |

<sup>1</sup> Within either PB, the expected percentage composition mirrors the global proportions

The percentage of pCONJ is higher among Common PTUs than Rare PTUs. Together, the proportion of pMOB and pOriT plasmids is also higher in the Common PB (40.4 %, CI_95%_ = [39.9, 41.0]) than in the Rare PB (37.0%, CI_95%_ = [36.4, 37.6]) (**Supplementary Table 3**). In contrast, pNT plasmids are more frequent in the Rare PB (44.4 %) than in the Common PB (23.6%). These results suggest that transfer ability plays a fundamental role in the success of Common PTUs, although 55.6% of the plasmids of the Rare PB are transferable, and 23.6% of the plasmids of the Common PB are non-transmissible.

## Are rare hosts receiving plasmids from phylogenetically close common hosts?

We have seen that several rare host species harbour plasmids from common PTUs. For example, there are 59 rare host species with plasmids only from the Common PB. How can we explain this observation? One possible explanation is that these hosts received the plasmid from phylogenetically close common hosts. That is likely the case of *Vibrio owensii*, a rare species with eight plasmids from three PTUs. *V. owensii* belongs to the genus Vibrio, which includes one common species and 53 rare ones, and to the family Vibrionaceae, which includes one common and 64 rare species. The common species within this genus and family shares all three PTUs with *Vibrio owensii*. However, this case of *Vibrio owensii* is far from general.

We have seen that, at least in the case of the *Vibrio* genus, a common host species harbouring plasmids from common PTUs likely contributes to the presence of these plasmids in rare host species of the same genus. To investigate the generality that rare hosts harbour common plasmids due to their taxonomic proximity to common hosts with the same PTUs, we performed the following analysis. For each of the 59 rare species, we looked at plasmids to check whether their PTUs were also found in common host species of the same genus or family. For 46 species (78%), there was no common host species within the same genus harbouring plasmids of any of the common PTUs found in the rare species. At the family level, 40 species (68%) had no common host species within the same family with plasmids of any of the same PTUs. As an example, we investigate the case of *Bacteroides cellulosilyticus*, a rare species, carrying three plasmids from two common PTUs: (i) one from a certain PTU and (ii) two plasmids from another PTU. In the first case, plasmids of the same PTU are also present in *Pseudomonas aeruginosa* and *Escherichia coli*, two common host species – both evolutionary distant from *Bacteroides cellulosilyticus* – sharing the same phylum, but not the same family, order, or class. This means that this PTU has a broad host-range (grade V ^12^). Therefore, this plasmid of a common PTU found in the *Bacteroides cellulosilyticus*, a rare host species, may have been transferred from *Escherichia coli* or *Pseudomonas aeruginosa*, two common hosts. In the case (ii), the two plasmids belong to a PTU that contains 20 plasmids in the dataset, 13 from the genus Bacteroides, six from the genus Parabacteroides, and one from the genus Phocaeicola, always from rare species. Hence, these two common plasmids found in the rare species *Bacteroides cellulosilyticus* do not appear to have been transferred from a common species (neither evolutionary close nor distant from *Bacteroides cellulosilyticus*).

## Discussion

It has been proposed that rare microbial species within ecosystems act as reservoirs of genetic resources, which may become accessible under favourable conditions, thereby potentially enhancing ecological stability through insurance effects ^1^. Our results show that, also in the case of plasmids, some taxonomic units are very abundant while many are very rare, and that rare PTUs harbour a strong diversity of genes, including virulence genes. The main caveat of this work is undoubtedly the fact that the plasmids analysed in this study were not derived from a single ecosystem, so caution is warranted when interpreting the broader implications of our findings ^10^. Yet, several findings concerning this dataset present striking parallels with important findings using data from ecosystems regarding plants, animals or microorganisms ^1,4,5,13^.

Recognizing the existence of rare biospheres within bacterial communities has been highly significant, as these bacterial species may constitute extensive reservoirs of genomic innovation ^3,5,21,22,22^. They have a great potential for dynamic adaptation because individual cells from low-abundance bacterial taxa can proliferate rapidly through clonal replication when environmental conditions shift to favour their growth ^4,5^. If this ability of clonal amplification is relevant for bacterial cells, it is even more so in the case of plasmids. These genetic elements may hitchhike the clonal expansion of their host, but they can also expand horizontally through bacterial communities, even between bacterial taxa, if the conditions are appropriate ^14,34,35^. Moreover, plasmids may even help other plasmids or be helped by other plasmids to transfer to other cells ^30,32,36–38^. These plasmids’ characteristics might annihilate the fat-tailed distributions frequently observed in bacterial communities, but our findings show that that was not the case.

The plasmids of this study have very diverse origins. Nevertheless, we observed the same striking distribution pattern as seen in microbial communities from a given ecosystem or habitat: characterized by a few highly abundant PTUs and a fat-tail of rare ones (**Figs. 1 a, b**). One possible explanation for this finding is that the PTUs have not been sampled exhaustively, a phenomenon referred to as the “veil line” in biodiversity studies. The veil line is an imaginary boundary that shifts progressively to the left of the distribution as sampling effort increases ^13^. As this shift occurs, a peak may emerge in intermediate abundance classes, rather than in the singleton class, as observed in the present data.

The fitting of the PTU Abundance Distribution (PAD) curve revealed that the Poisson log-normal distribution provided an excellent fit (**Figs. 1 c, d**). This result aligns with recent findings ^13^ and suggests a compelling similarity between plasmid abundance patterns and well-established results from other biodiversity studies. This observation is noteworthy, as the shape of the PAD may offer insight into the underlying processes of plasmid diversification, as well as the emergence and persistence of rare plasmids. If the PADs are genuinely monotonically decreasing, this would be a noteworthy result, as the PAD shape could provide valuable insight into the underlying processes of plasmid diversification, as well as the emergence and persistence of rare plasmids. For instance, under Hubbell’s Neutral Theory of Biodiversity and Biogeography ^24^, a monotonically decreasing function, such as the ones we observed, implies that new species arise with very few individuals. The PAD in our dataset peaks at singletons, likely reflecting a process of “speciation” in which novel PTUs emerge at initially very low abundance. This interpretation is consistent with the well-known ability of plasmids to change ^12,39–42^. These questions should be addressed in future research.

An important biological implication of the power-law is that some PTUs are highly successful (hence belonging to the Common PB). What is the cause of their success? If PTU’s success was attributed to that of their hosts, one should expect that all plasmids belonging to common PTUs should be hosted by common species and all plasmids belonging to rare PTUs should be hosted by rare species (0% mismatches). In contrast, if PTUs were randomly assigned as rare or common, 57% plasmids would not match the assignment of their hosts. We observed an intermediate number: 25% of the plasmids belong to a rare or a common species, despite belonging to a common or rare PTU, respectively. We conclude that some plasmids and the corresponding hosts share an evolutionary success. Yet, they are far from the majority. For example, the dataset includes 9,343 plasmids from *E. coli*—one of the most extensively studied bacterial species. Despite being the most common host, 1,649 of these plasmids (18%) belong to rare PTUs, and thus to the Rare PB. *Lactiplantibacillus plantarum*, the eighth most common host species, shows an even greater proportion: 77.4% of its plasmids (632 out of 816) belong to the Rare PB, with only 184 in the Common PB, or *Levilactobacillus brevis*, a common species with 104 of its 112 (92.86%) plasmids in the Rare PB. These examples also suggest that the success of plasmids from common PTUs is not necessarily driven by the success of their common hosts, nor are these hosts responsible for the success of their plasmids from the common PTUs. The same applies to plasmids from rare PTUs.

Intriguingly, the long list of rare PTUs is associated with exceptional diversity in both coding sequence clusters and virulence genes (**Fig. 3**). Therefore, gene diversity is not responsible for the success of the common PTUs. In contrast, the diversity of antibiotic resistance genes (ARGs) is higher among the plasmids from the common PTUs than in the rare ones. One possible explanation for such a contrast is that the sequencing of RefSeq genomes may have been motivated by antibiotic resistance. Yet, even if antibiotic resistance genes are less diverse in the Rare PB, 155 out of 761 genes (20.4%) are only present in the Rare PB. For example, resistance genes of the Nitroimidazole antibiotic class were found only in the Rare PB (**Supplementary Fig. 5**). Let us suppose that future studies confirm that most antibiotic resistance genes are indeed more often present in the Common PB. That might suggest that resistance genes in the Rare PB are not as relevant as, e.g., virulence genes. Yet this interpretation should be approached with caution. If new antibiotics are developed — particularly those with novel modes of action—potential resistance genes are more likely to emerge from CDS that have not yet been identified as ARGs, rather than from those already known. Our results, which reveal exceptionally high CDS cluster diversity among plasmids in the Rare PB, suggest that this group represents a vast gene pool from which novel resistance genes may evolve. As such, these findings serve as a warning: resistance to future antibiotics may arise from currently uncharacterized gene sequences within the Rare PB.

A recent study has shown that ARGs are preferentially acquired and carried by broad-host-range plasmids and that these plasmids are more ‘plastic’ (e.g., higher recombination rates or higher mobility) ^43^. We have found that one PTU belonging to the Rare PB is present in hosts from as many as four different phyla (**Supplementary Fig. 7**). Future studies should look for recombinases in these plasmids and study rates of allelic exchanges and changes of gene repertoires, as well as their mobility ^43^. This PTU is a peculiar case of an important characteristic of rare PTUs concerning host ranges: per number of plasmids in each PTU, the host range of rare PTUs is higher than common PTUs. Strikingly, we observed the same pattern at all taxonomic levels, from species to phylum (**Fig. 4** and **Supplementary Fig. 8**). One could argue that the overall broader host range of rare PTUs is irrelevant because these PTUs have very few plasmids. However, these PTUs are very numerous, representing 94% of the PTUs and comprising almost half of the plasmids in the dataset. Thus, these PTUs likely play critical roles in bacterial evolution when hosted by different prokaryotic species/genera/…/phyla, eventually co-inhabiting the same cell with other plasmids, from both plasmid biospheres.

In bacterial communities, it has been proposed that rare taxa tend to remain rare because they are less visible to predators, such as viruses or protists ^4,44^. Yet, our results strongly suggest that this explanation does not apply to plasmids. If it did, rare host species should exclusively harbour plasmids from the Rare PB, while common hosts should lack plasmids from this pool; this scenario is not supported by our data.

We have seen that the very good fit of a Poisson log-normal distribution to the PTU abundance distribution suggests that many new rare PTUs likely arise from genetic changes of the already existing pool of plasmids (from common or rare PTUs). Therefore, a potential mechanism underlying the emergence and/or persistence of plasmids in the Rare PB could be the acquisition of new genes, also likely to transform plasmids from a common PTU into a rare PTU. Nevertheless, this mechanism seems unlikely because plasmids belonging to Rare and Common PBs do not differ significantly in size (base-pairs), and plasmids in the Common PB contain only a slightly higher number of CDS (**Supplementary Fig. 4**).

Finally, plasmid mobility may explain the success of common PTUs. About 76.4% of the plasmids in the Common PB were transferable (pCONJ, pMOB, and pOriT plasmids), in contrast with the 55.7% transferable plasmids from the Rare PB. Consequently, the proportion of non-transmissible plasmids (pNT), is almost the double (44.4%) in the Rare PB than in the Common PB (23.6%) (**Table 2**). It is intriguing, however, that so many pNT plasmids are rare. There are some possible explanations, for example: (i) they may be misclassified as pNT, but future research may find *OriT*’s sequences on them ^14,45^; (ii) they belong to a PTU that also includes transferable plasmids; (iii) they were recently pOriT, pMOB or pCONJ, but they lost relevant transfer genes ^41,46^; or they easily transfer through alternative mechanisms, such as by transformation, or by integrating into the genomes of phages (transduction), or other plasmids (conduction) ^31,33^.

The dataset used in this study contained 20,580 genomes, 218 of archaea and 20,362 of bacteria, harbouring plasmids from 16,547 PTUs. Yet, none of these PTUs is present both in archaea and bacteria – even if one of the PTUs of archaea belong to the Common PB. This suggests that inter-domain plasmid transfers are rare events (possibly because their habitats and selective pressure do not coincide) ^47,48^.

Our study has important limitations. The first limitation was already mentioned throughout this study: plasmids (and their hosts, bacteria and archaea) studied here do not belong to a single ecosystem/ habitat. For example, *E. coli* is certainly not the most abundant bacterial species. Future studies should focus on plasmids from a single habitat. That would imply sequencing all plasmids present in a single microbiome. A second limitation concerns the dataset used in this study. The dataset is likely biased towards clinical bacteria. Different datasets have different characteristics, e.g. the number of plasmids carrying antibiotic resistance genes ^49^. A third limitation is the very low number of plasmids from archaea. Future studies should study their plasmids in this context. This limitation is related to the second one, as bacterial species have been of greater interest from the clinical or industrial viewpoints. Yet, despite these limitations, the excellent fit of a power-law to the rank abundance or the Poisson log-normal to the PTU abundance distribution is remarkable. That implies that PTUs are strongly unbalanced, leading to the definition of a rare and a common plasmid biosphere and the results shown here.

Our results strongly suggest that the rare plasmid biosphere may play critical roles in microbial ecosystems. Plasmids of rare PTUs act as reservoirs of diverse coding sequences, including genetic resources linked to virulence and antibiotic resistance, potentially providing an ‘insurance effect’ in response to selective pressures such as antibiotic use, as previously hypothesized for microorganisms ^1,50^. However, future research is needed to address key questions within specific habitats: (1) Do rare plasmid biospheres exist at the level of single habitats? (2) In which habitats or ecosystems are rare plasmid biospheres present? (3) Is the metabolic activity of cells harbouring rare or common plasmids different? (4) We have seen that plasmids of common PTUs have increased transfer ability; could other factors, such as encoding bacteriocins ^51^, be responsible for the success of the common PTUs?

Our finding opens new avenues in the quantitative biology of plasmids, in parallel with recent discoveries of universal laws relating plasmid size to protein-coding genes and copy-number ^52,53^. But this work has at least two main practical implications. First, researchers can carefully look at rare PTUs to identify possible candidate genes that confer resistance to antibiotics under clinical trials. Second, by studying plasmids of rare PTUs, we can also find relevant new genes, such as those involved in nutrient cycling, bioremediation, or pollutant degradation ^54–56^.

## Acknowledgments

João S. Rebelo and Célia P. F. Domingues acknowledge FCT-Fundação para a Ciência e a Tecnologia, IP for their fellowships (PhD grants SFRH/BD/04631/2021, and UI/BD/153078/2022, respectively). FCT also supports our research through the strategic funding to cE3c (DOI:10.54499/UIDB/00329/2020) and the associate lab CHANGE (LA/P/0121/2020). The authors thank Eduardo P.C. Rocha, Lounès Chichi and João. A. Gama for discussions and critical reading.

## Author contributions

Conceptualization, J.S.R., C.P.F.D., L.B.A., and F.D; Software, J.S.R., C.P.F.D. and L.B.A; Methodology, J.S.R., C.P.F.D., L.B.A., T.N., and F.D.; Investigation, J.S.R. and C.P.F.D.; Formal Analysis, J.S.R., C.P.F.D., L.B.A., T.N. and F.D.; Writing – Original Draft, J.S.R., C.P.F.D., L.B.A., and F.D.; Resources, T.N.; Visualization, J.S.R., C.P.F.D., and L.B.A; Supervision, T.N. and F.D.

## Methods

A total of 54,264 complete plasmids were retrieved from the NCBI RefSeq database (https://ftp.ncbi.nlm.nih.gov/genomes/refseq/), last accessed in February 2025^10^. To exclude potential secondary chromosomes, we removed 1,355 plasmids larger than 500 kb, resulting in a final dataset of 52,909 plasmids for our analysis.

We identified the putative coding sequences (CDS) using Prokka v1.14.6 ^57^. We then clustered the CDS using CD-HIT v4.8.1, with a 90% identity threshold. ARGs were identified using AMRFinderPlus v4.0.19 ^28^ (database version 2025-03-25), employing default parameters: sequence identity > 90%, e-value < 1e−20, and minimum coverage > 50%. VGs were detected using ABRicate v1.0.1 with a minimum identity of 90% against the Virulence Factor Database (VFDB) ^26^.

To cluster the plasmid sequences from the RefSeq database into Plasmid Taxonomic Units (PTUs) ^12^, we used the method described by Camargo *et al*. ^58^. This approach, similar to the one defined by Redondo-Salvo *et al*. ^12^, allows for the *de novo* grouping of plasmids based on average nucleotide identity (ANI) and aligned fraction (AF). As a result, all plasmids are assigned to a PTU, including those not previously associated with existing PTUs.

Sequence alignments were performed using blastn (parameters: –outfmt –6 std qlen slen’ –max_target_seqs 20000 –task megablast –evalue 1e-5). Afterwards, ANI and AF values were calculated using a Python script available at https://bitbucket.org/berkeleylab/checkv/src/master/scripts/anicalc.py. These values were used to compute the edges (*weight = AF × ANI*), which were subsequently applied to construct the Leiden graph ^59^ using the pyLeiden script available at https://github.com/apcamargo/pyleiden (parameters: ‘-n 5 – r 1.2’). Taxonomic information was retrieved from the NCBI Taxonomy database (https://www.ncbi.nlm.nih.gov/taxonomy), which provides curated taxonomic classifications maintained by the National Center for Biotechnology Information (NCBI) ^60^.

To detect conjugative systems, we used MacSyFinder (v2.1.3) ^61^ with the parameters “--models CONJScan/Plasmids all –-db-type ordered_replicon”. If a conjugative element (T4SS) was detected, the plasmid was classified as conjugative (pCONJ). If no conjugative element was found but a mobilisable element (MOB) was detected, the plasmid was classified as mobilisable (pMOB). Plasmids lacking both T4SS and MOB elements were further screened for an origin of transfer (*ori*T). Those containing an *ori*T were classified as pOriT, while the remaining plasmids were considered non-transmissible (pNT). To detect origins of transfer, we downloaded a database of 91 experimentally validated *ori*T sequences ^62^. A *blastn* (v2.14.0+) search was performed against this database using the parameters “-task blastn-short –evalue 0.01” ^63^. We filtered the results with identity and coverage higher than 80%.

## Statistics

The following statistical analyses were conducted in R v4.4.2 ^5^. The binomial test was used to calculate the percentage of each characteristic in the respective biosphere. Pearson’s chi-squared test, a non-parametric method, was applied to assess the dependence between two categories, with effect size measured using Cramer’s V (rcompanion package v2.4.36). Differences in mean values between two groups were tested using a t-test with unequal variance (Welch’s t-test), and effect size was calculated using Cohen’s d (effectsize package v1.0.0). To compare the prevalence of ARGs and VGs in each biosphere, we used odds ratio (epitools package v0.5.10.1). We used the nonparametric Mann-Whitney U test, also known as the Wilcoxon rank-sum test, to compare the means of two samples, and the effect size was calculated using the Glass rank biserial correlation coefficient (r) (effectsize package v1.0.0). Visualizations were performed with the ggplot2 package v3.5.1.

