## Supplemental File for "The Rare Plasmid Biosphere: A Hidden Reservoir of Genetic Diversity"

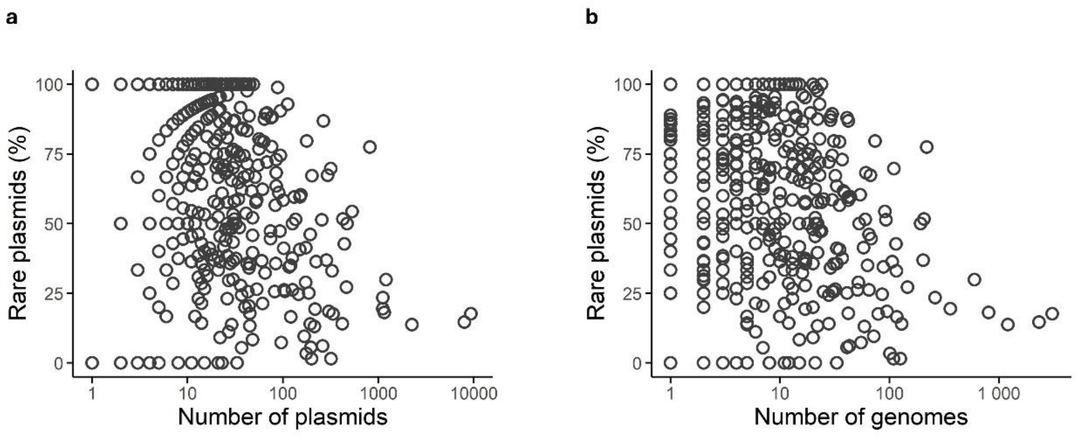


**Supplementary Fig. 1| The presence of plasmids from Rare PTUs across host genomes.** **a,** scatter plot between the number of plasmids in each species and the percentage of plasmids from the Rare PB. A moderate negative correlation was observed (Spearman's ρ = –0.484, $p < 2.2\text{x}10^{-16}$, n = 2363). **b,** scatter plot between the number of genomes for each species and the percentage of plasmids from the Rare PB. A moderate negative correlation was observed (Spearman's ρ = –0.485, $p < 2.2\text{x}10^{-16}$, n = 2363).

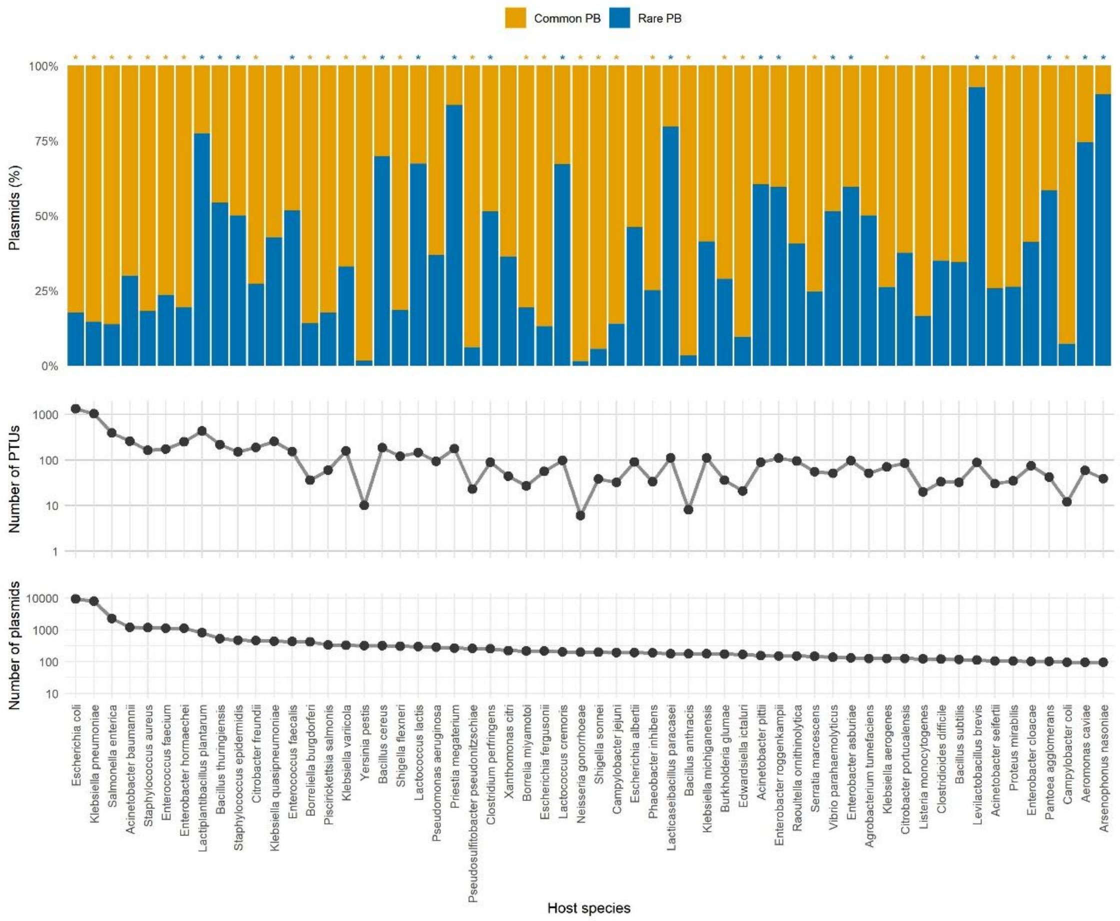


**Supplementary Fig. 2| Plasmids found in the common host species. a**, for each species, we see the percentage of plasmids in the Rare (blue) and Common (yellow) plasmid biospheres. Asterisks on the top represent species with significantly more plasmids of a plasmid biosphere than the other (binomial test, $p < 0.05$). **b,** number of different PTUs in each species. Note that a species may contain more than one plasmid of a PTU; in this case, we count them only once. Vertical axis in $Log_{10}$. **c,** number of plasmids in each host species. Vertical axis in $Log_{10}$.

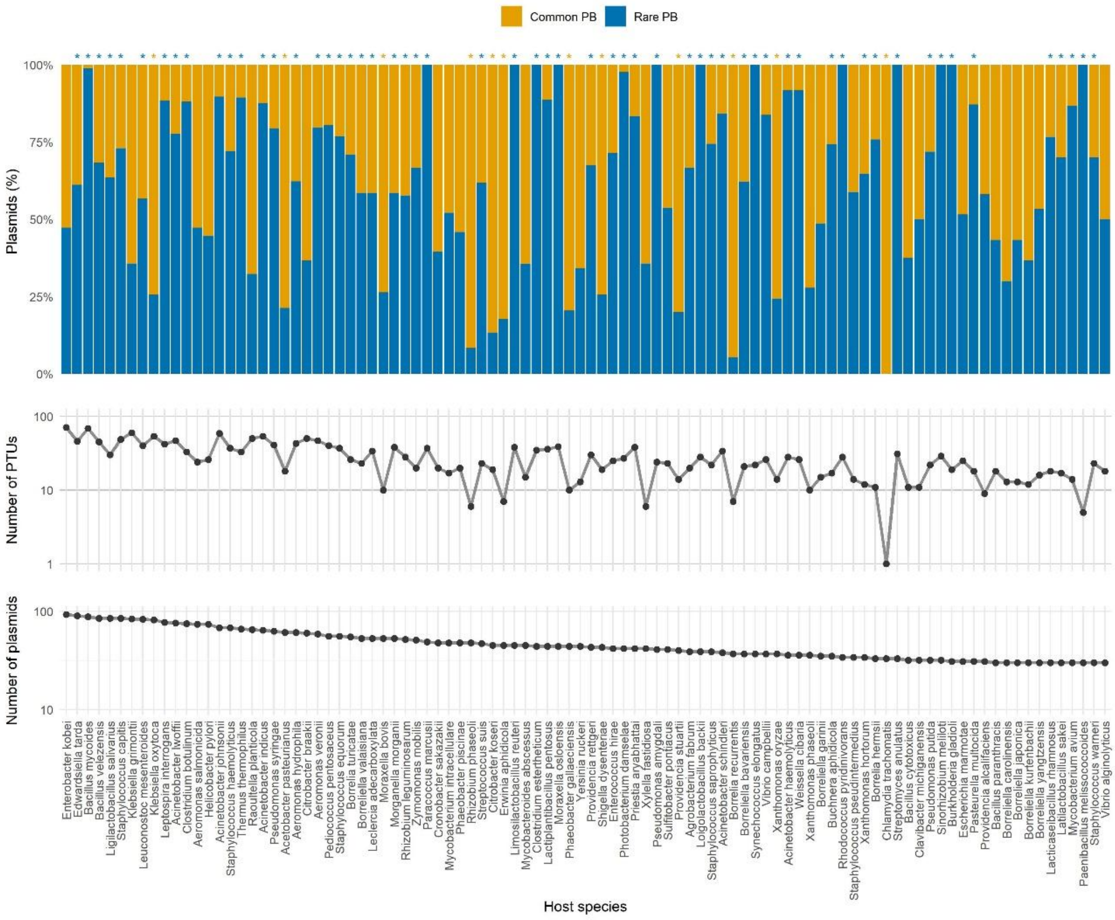


**Supplementary Fig. 3| Plasmids found in the rare host species. a**, for each species, we see the percentage of plasmids in the Rare (blue) and Common (yellow) plasmid biospheres. Asterisks on the top represent species with significantly more plasmids of a plasmid biosphere than the other (binomial test, $p < 0.05$). **b,** number of different PTUs in each species. Note that a species may contain more than one plasmid of a PTU; in this case, we count them only once. Vertical axis in $Log_{10}$. **c,** number of plasmids in each host species. Vertical axis in $Log_{10}$.

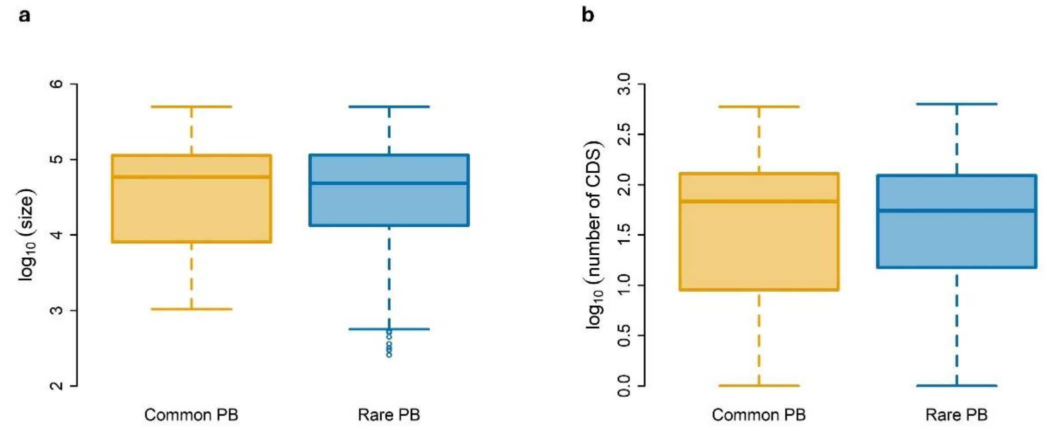


**Supplementary Fig. 4| Comparison of plasmid size in the two biospheres. a**, the number of base-pairs of plasmids in the Rare (blue) and Common (yellow) PBs are not statistically different (Wilcoxon: W = 341014327, *p* = 0.560; rank biserial = -0.003, $CI_{95\%}$ = [-0.01, 0.01]). **b**, the number of CDS is slightly lower in the Rare PB (blue) than in the Common PB (yellow) (Wilcoxon: W = 348118293, *p* = 0.0005), but with a tiny effect size (rank biserial = 0.02, $CI_{95\%}$ = [0.01, 0.03]).

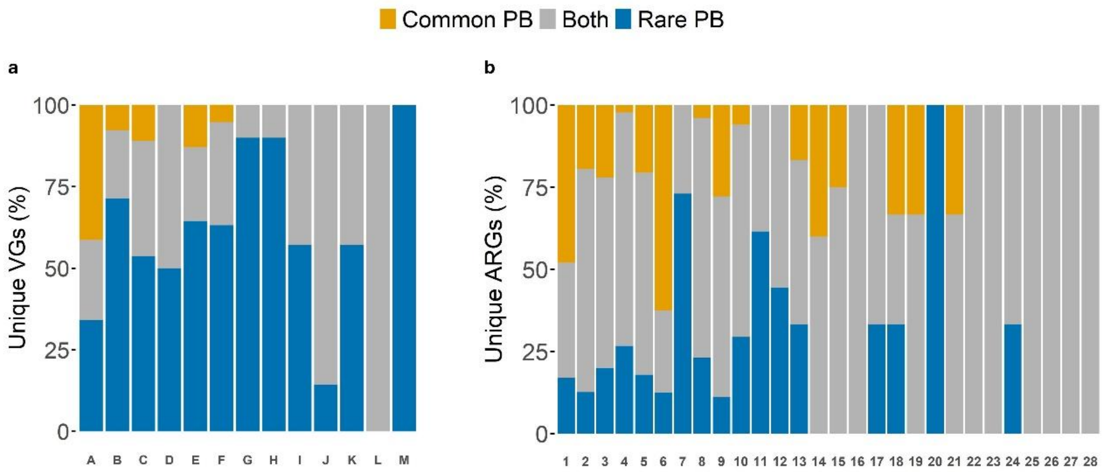


**A.** Effector delivery system
**B.** Adherence
**C.** Exotoxin
**D.** Nutritional / Metabolic factor
**E.** Immune modulation
**F.** Biofilm
**G.** Invasion
**H.** Motility
**I.** Exoenzyme
**J.** Others
**K.** Regulation
**L.** Antimicrobial activity / Competitive advantage
**M.** Stress survival

**1.** Beta-lactam
**2.** Aminoglycoside
**3.** Trimethoprim
**4.**Tetracycline
**5.** Quinolone
**6.** Colistin
**7.** Glycopeptide
**8.** Phenicol
**9.** Fosfomycin
**10.** Macrolide
**11.** Lincosamide
**12.** Lincosamide / Macrolide / Streptogramin
**13.** Sulfonamide
**14.** Phenicol / Quinolone
**15.** Aminoglycoside / Quinolone
**16.** Macrolide / Streptogramin
**17.** Bleomycin
**18.** Lincosamide / Macrolide
**19.** Lincosamide / Streptogramin
**20.** Nitroimidazole
**21.** Phenicol / Lincosamide / Oxazolidinone / Pleuromutilin / Streptogramin
**22.** Phenicol / Oxazolidinone
**23.** Rifamycin
**24.** Streptothricin
**25.** Avilamycin
**26.** Fusidic acid
**27.** Mupirocin
**28.** Streptogramin

**Supplementary Fig. 5| Presence of distinct virulence and antibiotic resistance genes per category.** Each bar represents the percentage of distinct virulence or antibiotic resistance genes in the Rare PB (blue), in the Common PB (yellow) or in both plasmid biospheres (grey). **a**, virulence genes **b**, antibiotic resistance genes.

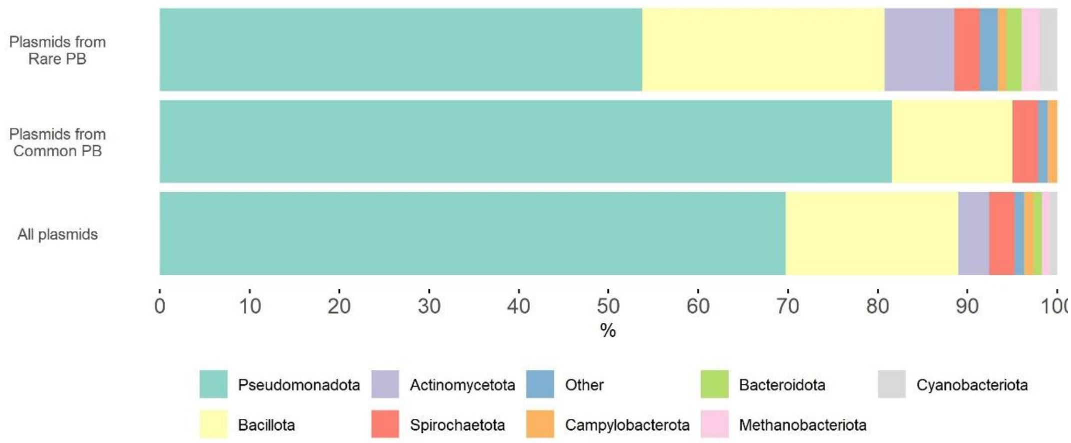


**Supplementary Fig. 6| Distribution of plasmids among hosts' taxa.** The most represented phyla of hosts harboring plasmids from the Rare PB, Common PB, and together. The most represented phylum for each plasmid biosphere is the same: Pseudomonadota, hosting 81.6% of the plasmids of the Common PB and 53.7% of the Rare PB. The 36,898 plasmids hosted by Pseudomonadota are classified in 8,784 PTUs – 688 PTUs from the Common PB and 8,096 PTUs from the Rare PB. The second most represented host phylum for each plasmid biosphere is also the same: Bacillota, hosting 13.4% plasmids of the Common PB and 27.0% of the Rare PB. The 10,152 plasmids hosted by Bacillota are distributed in 4,258 PTUs – 208 PTUs in the Common PB and 4,050 PTUs in the Rare PB. Species of these two phyla host 95.0% of the plasmids of the Common PB and 80.7% of the plasmids of the Rare PB. The third most common host phylum is not the same for the plasmids belonging to the Common PB (Spirochaedota, 2.8%) and Rare PB (Actinomycedota, 7.8%). The phylum Spirochaedota (2.8%) is in fourth place for the Rare PB. In general, plasmids from the Rare PB were found across more phyla than plasmids from the Common PB.

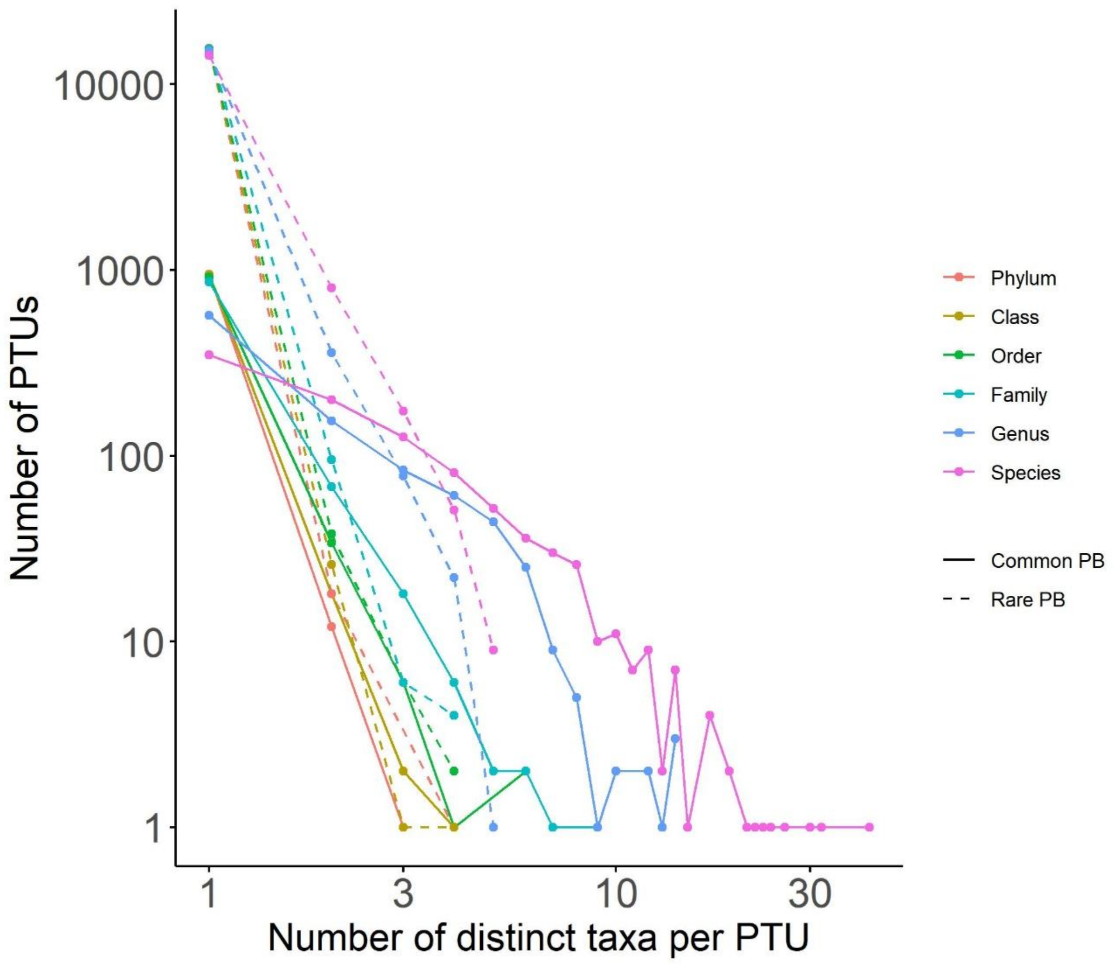


**Supplementary Fig. 7| Distribution of plasmids among hosts' taxa. T**he number of PTUs with a certain number of distinct taxa. Full lines: PTUs of the Common PB; broken lines: PTUs of the Rare PB.

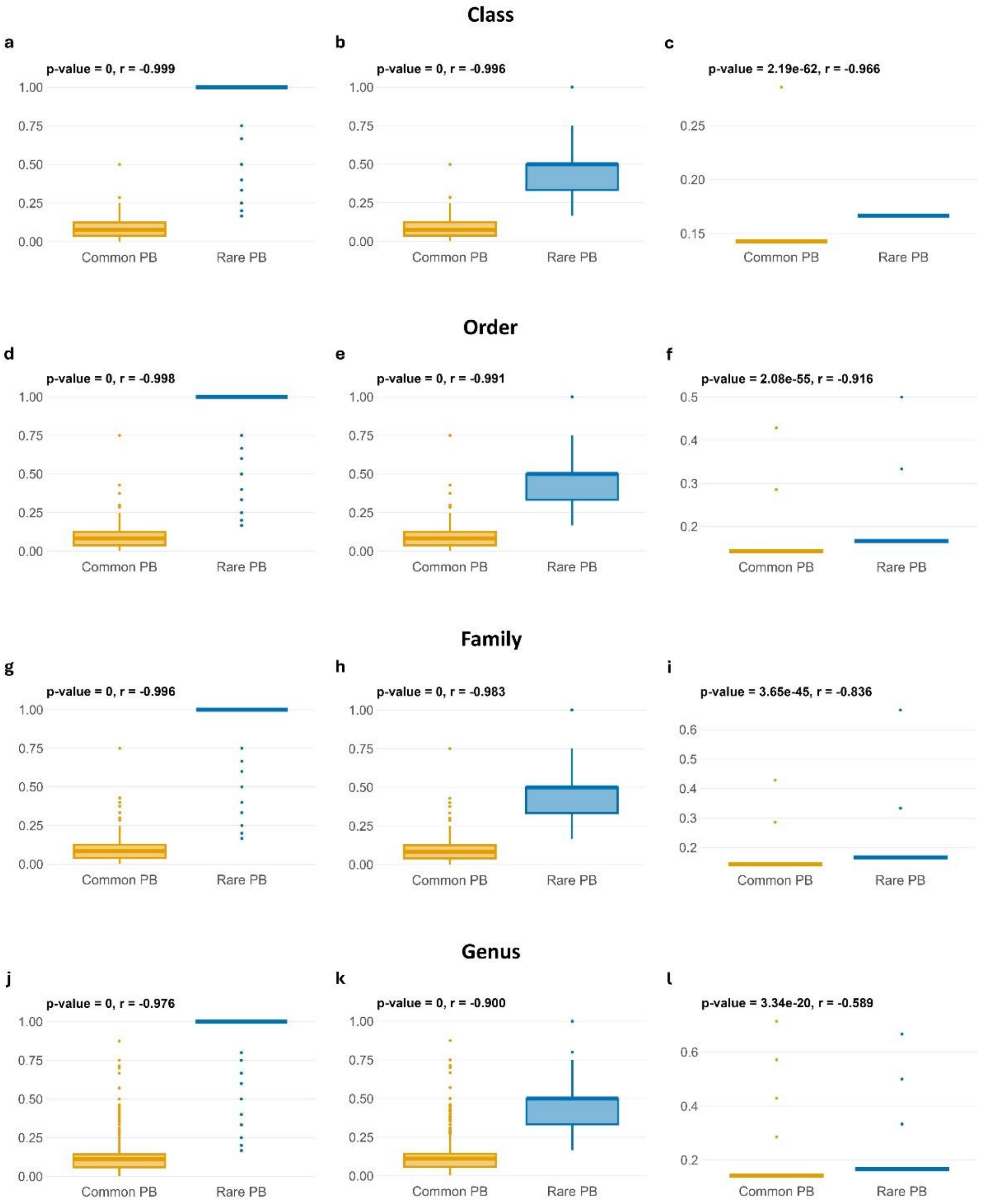


**Supplementary Fig. 8| The relative success of PTUs of the Rare and Common PB at the Class, Order, Family, and Genus levels.** The relative success of each PTU, measured as the number of hosts' taxa per number of plasmids in that PTU, is higher for the PTUs of the Rare PB (blue) than of the Common PB (yellow). This is the case for all levels, from the species to the phyla levels, including Genus, Family, Order and Class levels as shown here (Wilcoxon signed-rank test $p < 0.05$, large effect size in the twelve cases).

**Table S1| Akaike information criterion (AIC) values for the models using the negative binomial and Poisson lognormal distributions.**

| Model | AIC | $AIC-AIC_{min}$ |
|---|---|---|
| Poisson log-normal | 40956.77 | 0 |
| Negative binomial | 51125.05 | 10168.28 |

**Table S2| Estimated parameters for the negative binomial and Poisson log-normal distributions obtained using maximum likelihood methods.**

| Model | Parameter | Mean | SE |
|---|---|---|---|
| Negative binomial | μ | $1.4x10^{-7}$ | $2.8x10^{-7}$ |
| Negative binomial | r (shape) | $2.1x10^{-8}$ | $4.3x10^{-8}$ |
| Poisson log-normal | μ | -12.866 | 0.05 |
| Poisson log-normal | σ | 3.98 | 0.012 |
| Poisson log-normal | μ (linear) | 0.007 | $4.83x10^{-4}$ |
| Poisson log-normal | σ (linear) | 384.224 | 80.647 |

**Table S3| Plasmids' transfer ability in the Common and Rare PBs.**

| | | N | Obs[%] | $CI_{95\%}$ | E[%][1] | $CI_{95\%}$ |
|---|---|---|---|---|---|---|
| Common PB | pCONJ | 10928 | 35.9 | [35.4, 36.4] | 28.5 | [28.0, 29.0] |
| | pMOB+pOriT | 12309 | 40.4 | [39.9, 41.0] | 39.0 | [38.4, 39.5] |
| | pNT | 7191 | 23.6 | [23.1, 24.1] | 32.5 | [31.9, 33.0] |
| Rare PB | pCONJ | 4174 | 18.6 | [18.1, 19.1] | 28.5 | [27.9, 29.1] |
| | pMOB+pOriT | 8320 | 37.0 | [36.4, 37.6] | 39.0 | [38.3, 39.6] |
| | pNT | 9987 | 44.4 | [43.8, 45.1] | 32.5 | [31.8, 33.1] |
